# Modelling Auditory Enhancement: Efferent Control of Cochlear Gain can Explain Level Dependence and Effects of Hearing Loss

**DOI:** 10.1101/2025.11.10.687695

**Authors:** Swapna Agarwalla, Afagh Farhadi, Laurel H. Carney

**Author notes:** Indicates equal contribution (Swapna Agarwalla, Afagh Farhadi).

## Abstract

The role of medial olivocochlear (MOC) efferent gain control in auditory enhancement (AE) was investigated using a subcortical auditory model. AE refers to the influence of a precursor on detectability of targets. The absence (or presence) of a precursor component at the target frequency enhances (or suppresses) detection under simultaneous masking conditions. Furthermore, the enhanced target under simultaneous masking acts as a stronger forward masker for a delayed probe tone, known as AE under forward masking. Psychoacoustic studies of AE report findings that challenge conventional expectations, and the underlying mechanisms remain unclear. For instance, listeners with hearing impairment have AE under simultaneous masking but not forward masking (Kreft et al., 2018; Kreft and Oxenham, 2019), whereas listeners with normal hearing have level-dependent AE under forward masking (Kreft and Oxenham, 2019). Our model with MOC efferent gain control successfully replicated these findings. In contrast, a model without efferent gain control failed to capture these effects, supporting the hypothesis that MOC-mediated cochlear gain modulation may play a role in AE and its alteration by hearing loss.

## I. Introduction

Sensory perception is shaped by spatial and temporal context, which can enhance or suppress target stimuli. Contextual influences play a critical role in facilitating the detection of novel stimuli (Stilp et al. 2010) and preserving perceptual stability in dynamic environments (Kuhl, 1979). The phenomenon of auditory enhancement (AE) exemplifies a broader principle of contrast enhancement, wherein a target sound embedded within a complex background becomes perceptually more salient when the background without a component at the target frequency is presented as a precursor (Viemeister, 1980; Viemeister and Bacon, 1982; Carlyon, 1989; Byrne et al., 2011; Feng and Oxenham, 2015, 2018; Kreft et al., 2018; Kreft and Oxenham, 2019; Mehta et al. 2021).

Given that most natural sounds, including speech and music, occur in continuous sequences in which each sound influences the perception of the next, AE may support communication by dynamically adjusting auditory perception to changing acoustic environments. For instance, exposure to broadband noise with a sinusoidally rippled spectral envelope leads to the subsequent perception of white noise as having a complementary spectral ripple (Wilson, 1970). Similarly, when a harmonic complex is presented with a spectral envelope complementary to a vowel—containing valleys where formant peaks would normally be—a following complex of equal-amplitude harmonics is perceived as that vowel (Summerfield et al., 1984, 1987). Summerfield and Assmann (1989) demonstrated that AE facilitates concurrent vowel perception, supporting the segregation of multiple voices in complex listening environments. Stilp (2019) concluded that AE enables listeners to adapt to the spectral characteristics of a speaker’s voice, improving vowel identification across different talkers and acoustic conditions.

The current study focused on two common AE paradigms: AE under simultaneous masking (Fig. 1A, based on Kreft et al., 2018) and AE under forward masking (Fig. 1B, based on Kreft and Oxenham, 2019). We chose not to include frequency roving, which has been included in other AE paradigms (e.g., Viemeister, 1980; Kreft & Oxenham, 2019; Byrne et al., 2011; Kreft & Oxenham, 2023). Although AE can exceed 20 dB with frequency roving, it varies substantially across listeners.

**Figure 1:**
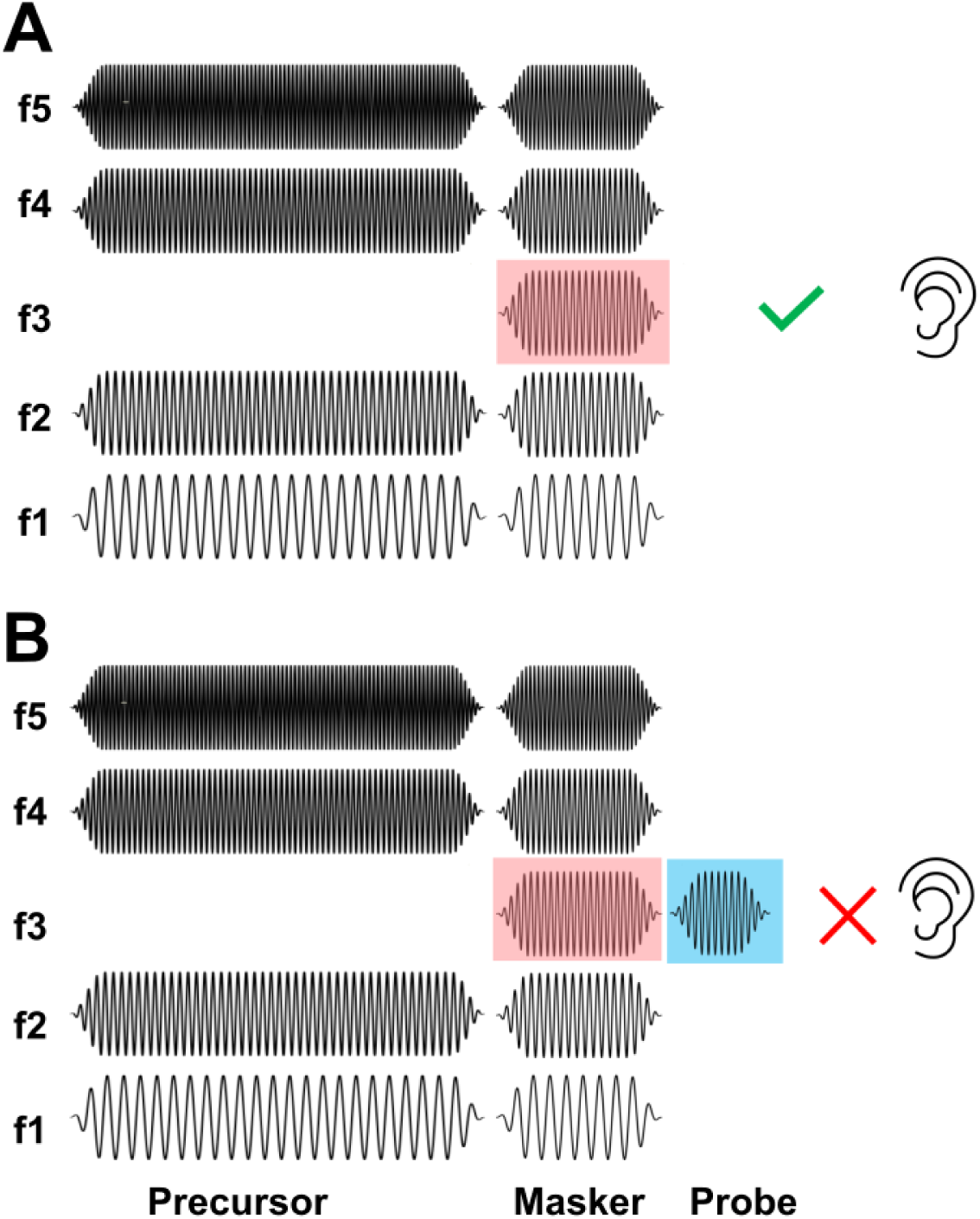
Schematic showing the paradigms of Auditory Enhancement (AE) used in the current study. A) AE under simultaneous masking: A precursor comprising the same background as the masker but without the target enhances the detectability of the target (red) making it “pop out” of the background. B) AE under forward masking: The precursor and masker are followed by a probe tone (blue). The target is a forward masker of the probe tone; when the target is enhanced, it is a more effective forward masker, increasing the probe-detection threshold.

Under simultaneous masking, the detectability of a target sound embedded in a background masker is enhanced when it is preceded by the background without the target-frequency component, making the target perceptually “pop out” (Viemeister, 1980; Carlyon et al. 1989; Wright et al. 1993; Kreft et al., 2018) (Fig. 1A, red). In contrast, under forward masking, the precursor and masker are followed by a probe tone (Fig. 1B, blue); the enhanced target (red) serves as a more effective forward masker of the probe, which leads to an increase in the probe-detection threshold compared to an unenhanced target (same as Fig. 1B, but without the precursor) (Viemeister and Bacon, 1982; Thibodeau et al. 1991; Wright el al. 1993; Byrne et al., 2011; Kreft and Oxenham, 2019).

One possible explanation for AE is that the precursor and masker have identical spectral characteristics and are perceived as a single auditory object, promoting segregation and enhancing detectability of the target component (Carlyon, 1989). Neural adaptation within individual frequency channels has been proposed as another potential mechanism, whereby adaptation to the precursor reduces the response to the masker, leading to a relative enhancement of the target (Delgutte, 1990; Oxenham and Plack, 1998). However, Viemeister and Bacon (1982) reported that the amount of forward masking produced by the target component increased when a precursor (which itself produced no forward masking) was added, suggesting an enhancement of the signal component which cannot be explained by neural adaptation. Additionally, neural adaptation does not explain why target loudness is enhanced by a precursor, whereas masker loudness remains unaffected (Wang and Oxenham, 2016). To address these limitations, an alternative hypothesis known as “adaptation of inhibition” has been proposed (Viemeister and Bacon, 1982). This hypothesis suggests that the components of the masker and precursor exert mutual inhibitory effects, which adapt over time. Over the course of the precursor, this inhibition adapts, leading to a stronger response to the subsequent target. Supporting evidence includes a report that the loudness of the target in the masker is enhanced when following a precursor (Wang and Oxenham, 2016).

Physiological studies have yet to establish a definitive neural basis for AE. Studies examining cochlear and neural contributions to AE have yielded mixed results. Physiological research in nonhuman species has found no evidence for AE at the auditory-nerve (AN) level in anesthetized guinea pigs (Palmer et al., 1995), yet single-unit recordings in awake marmosets have identified AE-related responses in the inferior colliculus (IC) (Nelson and Young, 2010). Similarly, human studies have failed to detect AE-related activity either in otoacoustic emissions (Beim et al., 2015) or auditory steady-state responses (Carcagno et al., 2014). Despite the extensive studies on AE, the underlying neural mechanisms remain unclear, particularly in the context of hearing loss. Also, the contribution of the efferent system, and its control of cochlear gain, to AE remains uncertain, largely due to the limitations of available physiological techniques. In particular, the difficulty in recording from AN fibers in unanesthetized animals limits our understanding of the influence of efferent gain control on peripheral coding because anesthesia suppresses efferent activity (Guitton et al., 2004).

A previous computational model with medial olivocochlear (MOC) efferents accounts for several auditory phenomena, including overshoot (Farhadi & Carney, 2023), forward-masking recovery (Maxwell et al., 2024), and recovery from forward masking (Agarwalla et al., 2025), phenomena that models without efferent pathways fail to explain. The current study investigated whether a subcortical model with efferent control of cochlear gain could explain AE. We hypothesized that the difference between MOC control of cochlear gain in response to targets with and without precursor stimuli explains AE. We replicated two psychoacoustic experiments: Kreft et al. (2018), which examined AE under simultaneous masking, and Kreft and Oxenham (2019), which investigated AE under forward masking, across different stimulus configurations and for listeners with and without hearing loss.

## II. Methods

### A. Models

AN responses were modeled with or without gain control by MOC efferents using the Farhadi et al. (2023) AN model. The Farhadi et al. model without efferent gain control is equivalent to the Zilany et al. (2014) AN model. As described in Farhadi et al. (2023), the model uses the LSR AN fiber response to represent the wide-dynamic-range inputs to the MOC system from the cochlear nucleus, some of which predominantly receive input from LSR and MSR fibers (Fekete et al., 1984; Ye et al., 2000; Romero and Trussell, 2021). The Farhadi et al. (2023) model also includes a fluctuation-driven signal from the IC, which reduces cochlear gain (Farhadi et al., 2023). The input to the MOC system from the IC is driven by HSR AN fibers. The two input signals are first scaled and summed, then passed through a first-order low-pass filter. The filtered signal was the input to a rational input-output function representing the MOC stage, which converts the input into a cochlear-gain factor (Farhadi, 2023). Specifically, the MOC output is a scalar between zero and one, the gain factor, which is multiplied by the output of the control path to adjust cochlear gain dynamically. The model also includes a physiologically based MOC time constant (τ_MOC_ = 260 ms) to limit the dynamics of efferent gain control (Farhadi et al., 2023). In the AN model, there is no explicit gain parameter; rather the peripheral-filter time constant is varied to manipulate gain and bandwidth (Patterson et al. 1988; Carney, 1993; Zilany and Bruce, 2006; Zilany et al., 2014); due to the gain-bandwidth trade-off, gain is proportional to the filter time constant. We plot the dynamic changes in the time constant below for both AE paradigms studied here; for the Farhadi et al. (2023) AN model, the change in cochlear gain is 2.5 times the change in the time constant (see Zilany and Bruce, 2006).

Tuning in the AN model was based on human frequency-selectivity data (Ibrahim and Bruce, 2010; Shera et al., 2002). The average audiograms for listeners with normal hearing (NH) and with hearing impairment (HI) in the psychoacoustic studies (Kreft et al. 2018; Kreft and Oxenham, 2019) were used to adjust the outer- and inner-hair-cell parameters of the AN model for each frequency channel (Zilany and Bruce, 2006), assuming that two thirds of the loss was due to outer-hair-cell impairment and one third due to inner-hair-cell impairment.

AN-model responses provided the inputs to a simple IC model for AM tuning (Mao et al., 2013), a band-pass modulation transfer function (MTF) with a best modulation frequency (BMF) of 64 Hz, the median BMF for IC cells that are excited by modulations, as reported by Kim (2020). Band-enhanced (BE) neurons, which exhibit elevated discharge rates over a range of modulation frequencies (Kim et al., 2020) were the only IC cell type modelled in this study. Thresholds for detection of probe tones were estimated using discharge rates from a same-frequency inhibition-excitation (SFIE) model for IC neurons (Nelson and Carney, 2004). The input to the SFIE model was from high-spontaneous-rate AN fibers from the AN model with or without efferent gain control.

## III. Model predictions of psychoacoustic experiments

### A. Experiment 1: Auditory enhancement under simultaneous masking

#### 1. Stimuli

The stimuli had a 500-ms-duration precursor followed by a 20-ms gap and then a 100-ms-duration masker. Maskers and precursors were gated on/off with 10-ms cos^2^ ramps. Maskers had four equal-amplitude, logarithmically spaced frequency components (2462.3, 2639.0, 6062.9, and 6498.0 Hz) in sine phase, centered around the target frequency of 4000 Hz. The precursor was a copy of the masker with or without the target component. Three stimulus conditions were simulated (Kreft et al., 2018, Fig. 2):

**Figure 2:**
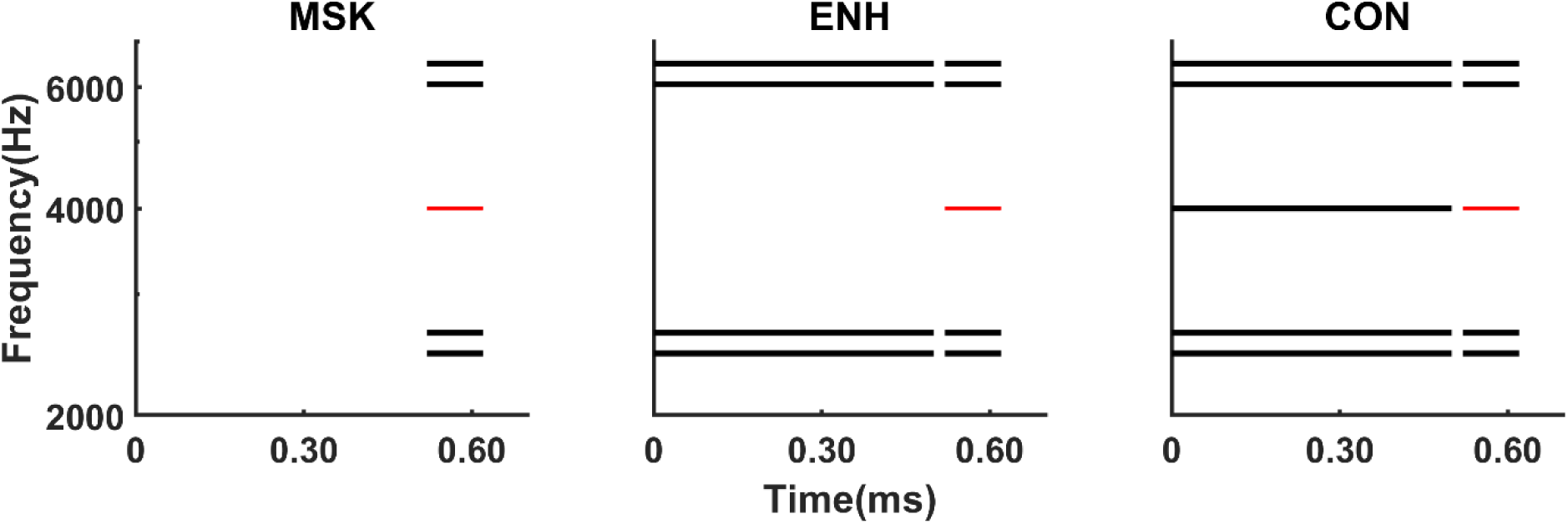
Schematic diagrams of stimuli from Kreft et al. (2018) used for AE under simultaneous masking stimulus conditions. The masker-only condition (MSK) included the masker (black lines) and the target (red). The enhanced condition (ENH) included a precursor without the target component, enhancing the target perception. In contrast, the control condition (CON) included a precursor with the target component. The simulations used a two-interval, two-alternative forced-choice task; only the intervals that contained the target are shown.

MSK = Masker with target component (Control condition 1)

ENH= Added precursor without target component (Enhanced condition)

CON = Control; Added precursor with target component (Control condition 2)

For the NH model, simulations were performed for two level configurations:

(1) NHSL: Masker components were 20 dB above the average absolute probe-detection threshold at the target frequency (4000 Hz) for listeners with normal hearing (37 dB SPL/component, from Kreft et al., 2018), and (2) NHSPL: Masker components were 20 dB above the average threshold at the target frequency (4000 Hz) in the presence of threshold equalizing noise (TEN, 52dB SPL) (Moore 2000). The target level in TEN noise was equal to the average SPL used for the listeners with HI in Kreft et al. (2018). TEN noise was gated on 100 ms prior to the start of the first interval and gated off 100 ms after the end of the last interval in each trial.

#### 2. Anticipated Results

For the ENH condition, the precursor would reduce cochlear gain in the off-target channels, but not in the target channel, thus the target component would elicit a relatively strong response. The relative advantage, or enhancement, of the target would be missing in the MSK condition, which has no precursor, and in the CON condition, for which the precursor contains the target frequency. The magnitude of enhancement would be reduced for HI listeners and for NH listeners in the presence of TEN noise due to overall reduced cochlear gain.

#### 3. Procedure

Model thresholds for probe detection were estimated using a two-interval, two-alternative, forced-choice, method-of-constant-stimuli paradigm with an inter-stimulus interval (ISI) of 500-ms. A total of 50 trials were simulated for each stimulus condition; half of the trials had the probe in the first interval and half had the probe in the second interval. Probe level was varied to estimate detection thresholds. Thresholds were estimated for the model with or without hearing loss for all stimulus conditions. Probe level was varied from 10 to 100 dB SPL, with 5 dB increments. For NHSL, NHSPL, and HI conditions, simulations were done at 37, 72, and 72 dB SPL per component level, respectively. For NHSPL, the TEN noise level was set to 52 dB SPL, to match the NH model threshold for detection of the probe alone to the mean threshold for the listeners with HI in Kreft et al. (2018). The levels used for simulations were matched to the model thresholds obtained by matching the approach for the psychoacoustic study by Kreft et al. (2018).

AN and IC responses were simulated for 41 channels with log-spaced characteristic frequencies (CFs) spanning 2-octaves, in steps of 0.1 octave, centered at 4000 Hz for Exp.1. Profiles of the model average-rate responses across CF channels (Fig. 3) were used to compute a decision variable for each interval within each trial. The presence of the target component had to be detected amidst the responses to the background masker. Average IC model discharge rate profiles across CF were computed over the time window that included the masker. Rates for CFs near the target frequency (Fig. 3, gray) were scaled by a factor K as a simple mechanism for differential cue weighting, assuming that on-frequency channels are given more weight than off-frequency channels. After scaling, the interval with the maximum value in the IC-BE rate profile was selected as the target interval (Fig. 3, asterisks). The percent correct was computed for each probe level, and a logistic function was then fit to each percent-correct vs. level curve. Threshold was estimated as the level for which the curve intersected the 70.7%-correct point. The scaling factor, K, was set to 2.5 by fitting the model threshold to the psychoacoustical threshold for MSK NHSL (as in Zaar and Carney, 2022). The value of K was then held constant for all other conditions.

**Figure 3:**
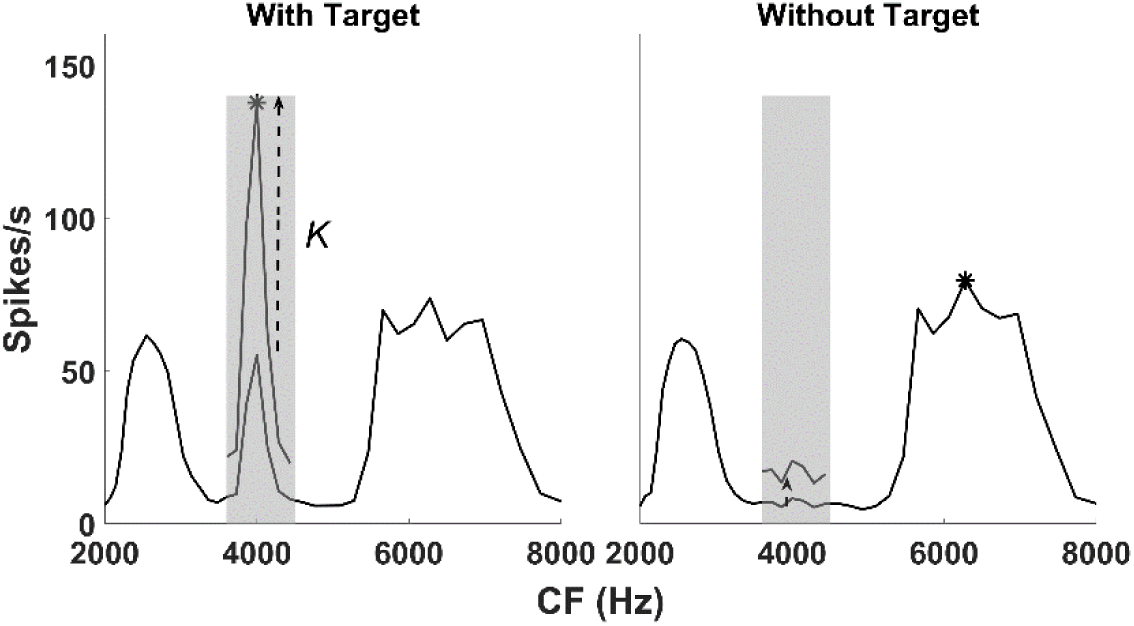
Decision variable for AE under simultaneous masking paradigm. Response profile as a function of CF at the IC stage of the model, showing the maximum response within the analysis window corresponding to the masker duration. The response profile is shown for a two-interval, two-alternative forced-choice task simulation, with (left) and without (right) the target 5 dB above the threshold. The narrow frequency band around the target frequency in the masker is highlighted in the gray shaded area and is scaled by a factor K, which was set to 2.5 to fit the model threshold to the psychoacoustic threshold for MSK NHSL. The interval with the greater maximum response was identified as the trial containing the target. The maximum response for the with-target and without-target intervals above are indicated by asterisks.

#### 4. Results

Model-AN rate profiles for frequency channels matched to stimulus components in response to the masker with a target (5 dB above absolute target threshold), illustrate the effect of efferent gain control on the simulations for NH listeners (Fig. 4). The stimuli were presented in quiet at a sensation level comparable to that for listeners with HI (NHSL; Fig. 4). Fluctuations in the AN response at the target frequency resulted from beating between low-frequency components (2639-2462=177 Hz). Without efferent input, model responses during the masker were similar across conditions (Fig. 4, top), but with efferent control, responses varied (Fig. 4, bottom). During the masker, off-target channels showed high onset responses for the MSK condition due to high peripheral-filter time constants (Fig. 5A, blue) and thus high cochlear gain. In the target channel (4000 Hz), cochlear-gain reduction was similar for MSK (blue) and ENH (red) but was further reduced for CON (yellow) during the masker, leading to lower AN rate responses. In HI listeners (Fig. 5B) and NH listeners with TEN noise (NHSPL, Fig. 5C), peripheral-filter time constants, and thus cochlear gain, was reduced overall compared to NHSL. The differences observed at the level of the AN were further accentuated at the level of IC.

**Figure 4:**
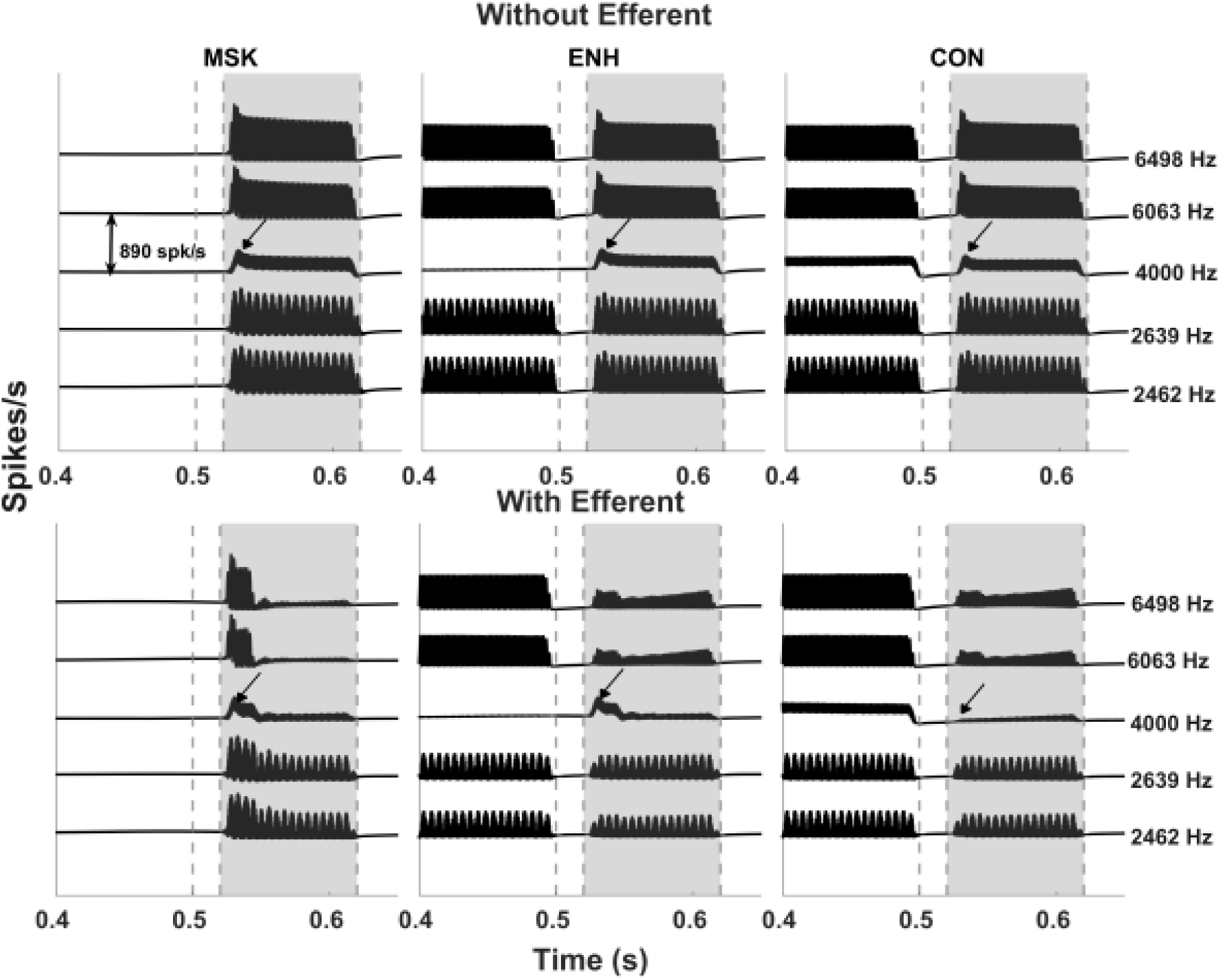
Efferent gain control in model results in differences in populations responses at the level of AN for AE under simultaneous masking under different conditions MSK (control 1), ENH (enhanced condition) and CON (control 2). The model without efferents showed subtle differences across stimulus conditions (top row) for the NH model in quiet (NHSL) for masker response (gray area). The model with efferents had lower responses in the off-target channels for ENH compared to MSK, whereas for CON the response in the target-frequency channel was lower than for both ENH and MSK (arrows). The dashed vertical lines in gray indicate the precursor offset, masker onset and masker offset respectively. The target is set 5 dB above the threshold observed for ENH condition.

**Figure 5:**
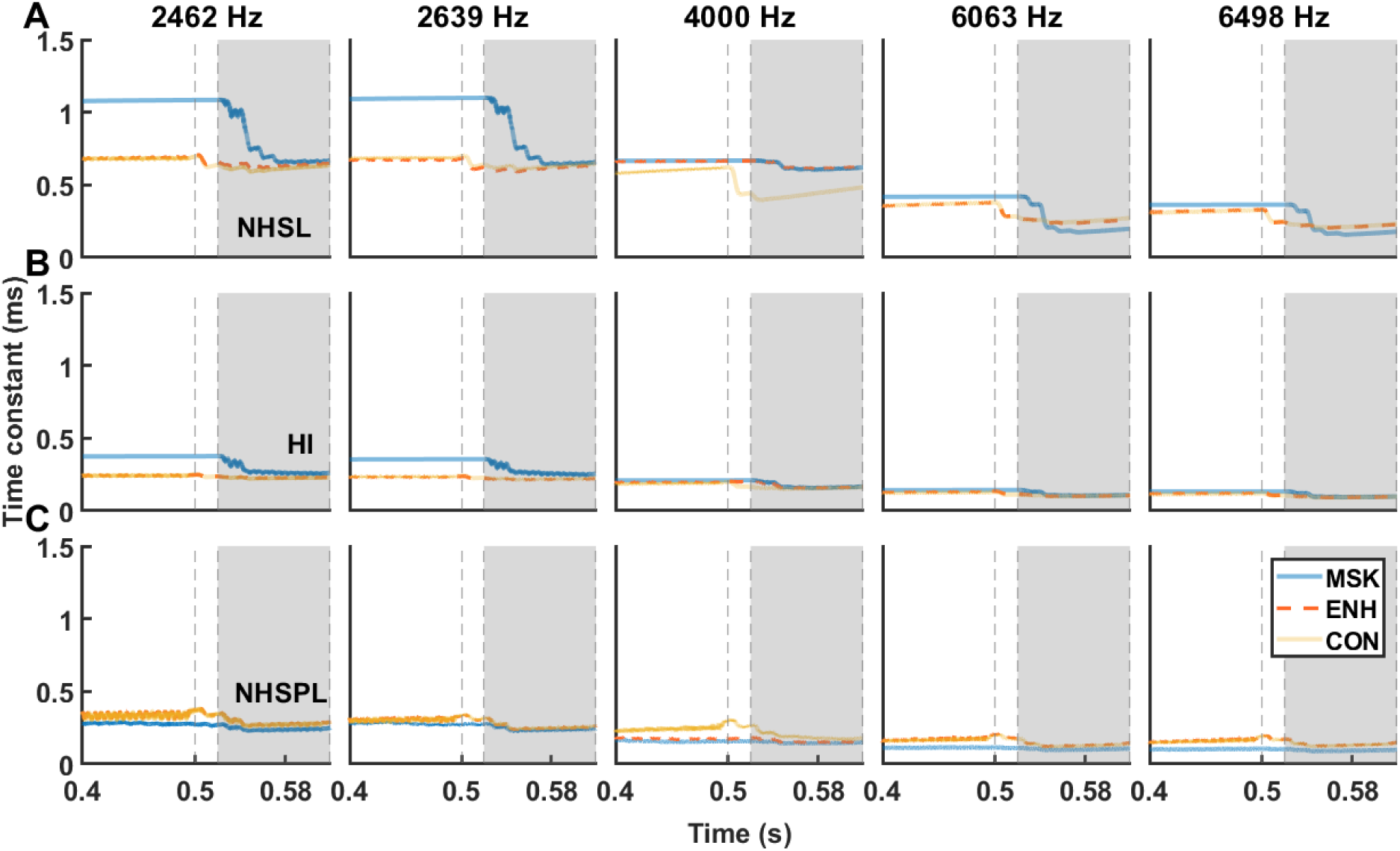
Changes in the time constant of the peripheral filter (the C1 filter in Zilany et al., 2014) during AE under simultaneous masking, for frequency channels tuned to stimulus components. There is no difference in the time constant at the target frequency (4000 Hz) during the masker (gray shaded area) for MSK (unenhanced, blue) and ENH (enhanced, dashed red) condition; the red and blue lines are overlapping. The dynamics of the time constant, and thus cochlear gain (see text) are shown for MSK (blue), ENH (red) and CON (yellow) conditions for A) NHSL, B) HI, and C) NHSPL configurations. The target was 5 dB above the threshold for the ENH condition.

Model IC rate profiles in response to the masker, with and without a target (5 dB above absolute target threshold), illustrate the influence of efferent gain control on the IC simulations (Fig. 6). For the model with efferent input (left, Fig. 6A), in the ENH condition the NHSL had a higher response rate in a narrow CF range around the target frequency (vertical line) when the target frequency was present compared to when it was absent. In contrast, for the MSK condition, the off-CF response dominated the narrow range around the target-CF channel (Fig. 6, left), making the task more challenging and resulting in higher detection thresholds for MSK compared to ENH for the same masker and probe level.

**Figure 6:**
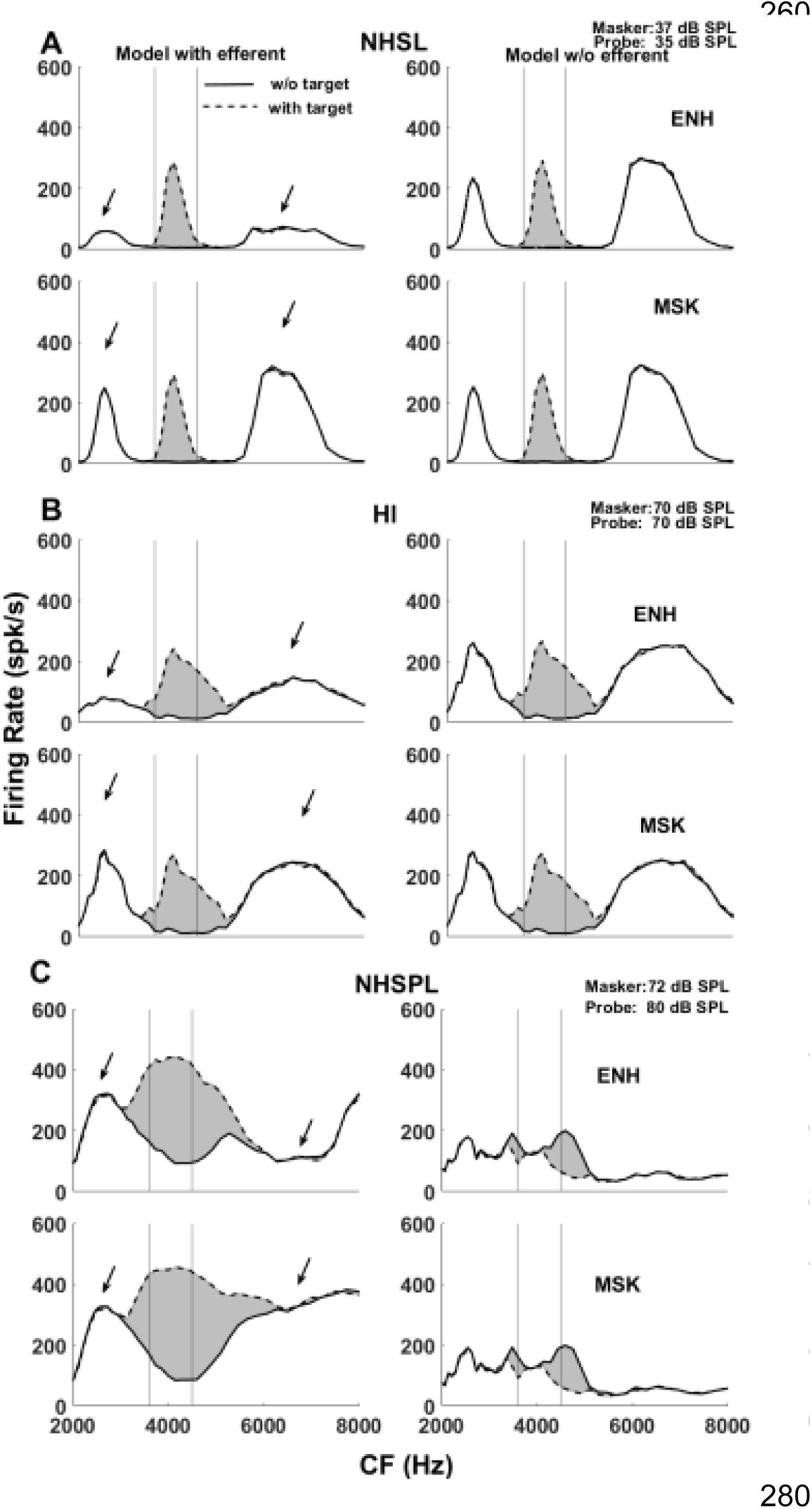
Efferent gain control in the model results in differences in population responses that are consistent with enhancement (left column). The model without efferents shows little differences across stimulus conditions (right column.) Response profiles as a function of CF are shown with (solid line) and without the target frequency (dashed line) for ENH (top) and MSK (bottom). The shaded gray region represents the area between the with- and without-target frequency, for the model with efferents on the left and without efferents on the right. The target was 5 dB above the threshold for the ENH condition. (A) NHSL: The NH model with stimuli presented in quiet at a sensation level (SL) comparable to the HI model. (B) HI model response. (C) NHSPL: NH model with stimuli presented in a background of TEN to simulate the hearing loss of the matched HI listener, with masker and enhancer presented at the same SPL as for the HI model. The two vertical lines in gray indicate the narrow range around CF which is compared to the off-CF channels on either side.

For the model without efferent gain control (Fig. 6A, right) there were subtle differences between firing rates in the MSK and ENH conditions due to stronger neural adaptation in the ENH condition. Efferent effects were minimal in the MSK condition for the following reason: the decision variable was the maximum response during the masker interval, typically corresponding to the onset response. The relatively slow efferent effects did not affect the onset responses, thus the thresholds of the models with and without efferents were similar (Fig. 6A).

In the HI model with efferent input (left, Fig. 6B), response profiles were broader, and had lower firing rates near the target frequency but higher rates at frequencies above the target, compared to the NHSL (Fig. 6A) which would lead to less enhancement in HI compared to NHSL. Thresholds were lower for the ENH condition than for MSK, similar to the trend in thresholds observed for NH listeners. In the model without efferent gain control, response profiles for MSK and ENH remained similar.

For the NHSPL configuration, thresholds were lower for ENH than for MSK. However, due to the presence of TEN noise, the response profiles differed from those in quiet (Fig. 6C); the tuning was broader compared to HI, although the stimulus level per component was similar for HI and NHSPL configurations. In the model without efferent gain control, the response in the without-target condition surpassed that in the with-target condition. This discrepancy arose because adding the target frequency to the TEN noise reduced the fluctuations in AN responses, resulting in a reduction of IC-BE responses, unlike other conditions for which responses increased with addition of the target. Consequently, the model without efferent gain control failed to reach the detection threshold in the NHSPL configuration, as the decision variable relied on the maximum response across trials, leading to an incorrect classification of the target-present condition. In NHSPL, thresholds for the models with and without efferent gain control differed for the MSK condition (Fig. 6C), unlike for the NHSL (Fig. 6A) and HI (Fig. 6B) conditions. The continuous presence of TEN noise in NHSPL, starting 100 ms before the stimulus onset, broadened the tuning in the NH model, explaining the differences in observed threshold with and without efferents, which were due to both compression and cochlear-gain reduction by the TEN noise throughout the stimulus. The overall pattern of results was consistent across all listener groups, with thresholds being highest in the CON condition, lower in the MSK condition, and lowest in the ENH condition. The thresholds for the model with efferent feedback (orange circles) closely matched the experimental data (violet squares), whereas the model without efferents (unfilled gray circles) were similar across most conditions (Fig. 7).

**Figure 7:**
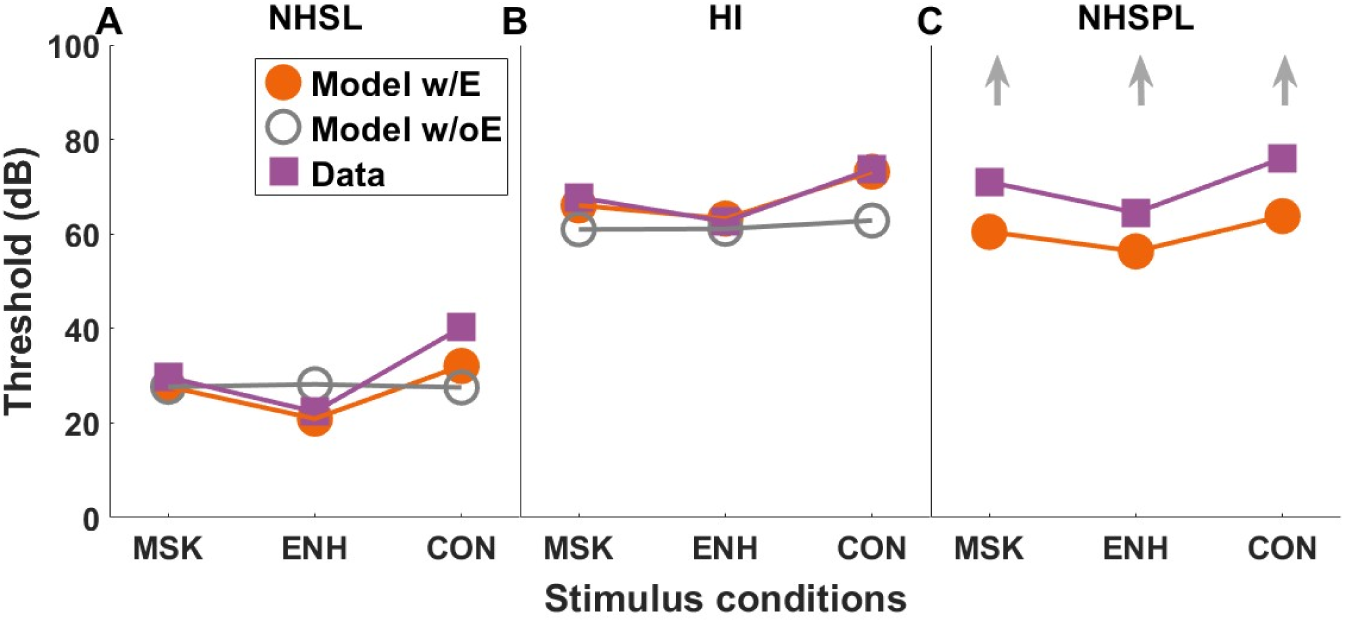
Thresholds for each stimulus condition for a target frequency of 4000 Hz across different configurations: (A) NHSL (B) HI (C) NHSPL. Data from Kreft et al. (2018) are shown as violet squares, model with efferent (w/E) as orange filled circles, and model without efferent (wo/E) as gray unfilled circles. Upward arrows indicate thresholds that were not reached.

The amount of enhancement was quantified for each configuration using the difference (in dB) between MSK and ENH thresholds (Fig. 8). Notably, the model with efferent feedback exhibited enhancement comparable to the psychoacoustic data, highlighting the possible role of efferents in AE. Note that the model without efferent feedback failed to reach threshold for the NHSPL configuration with TEN noise (upward arrows).

**Figure 8:**
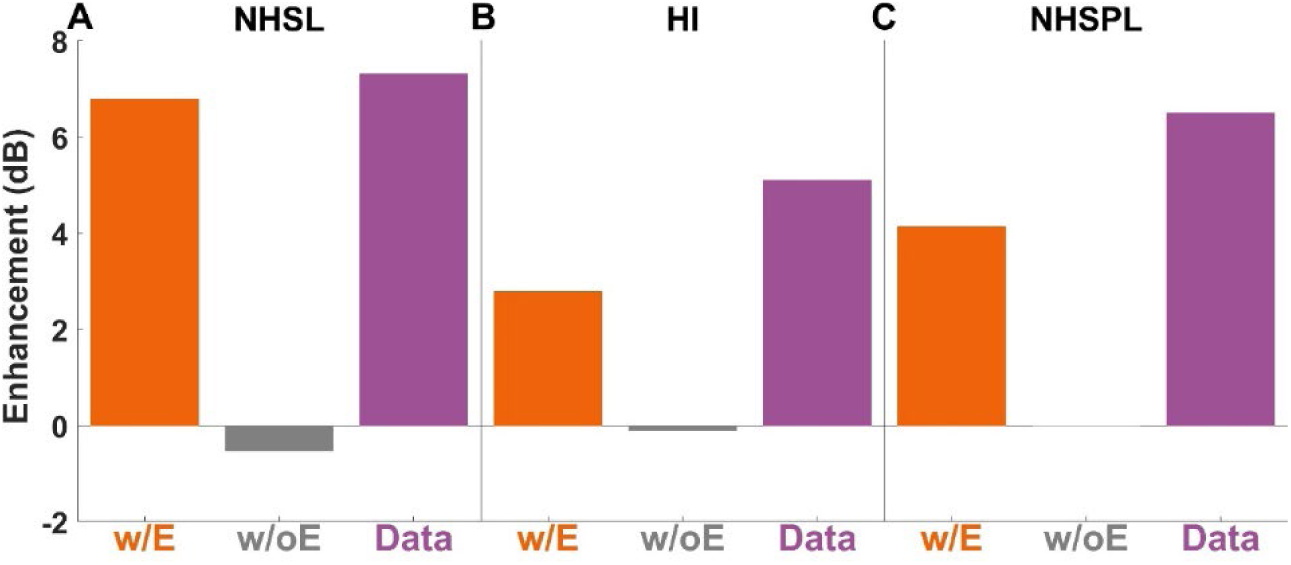
Enhancement (in dB) was computed as the difference between MSK and ENH thresholds. (A) NHSL, (B) HI, and (C) NHSPL configurations. Results are shown for the model with efferent (orange bar), model without efferent (gray bar), and data from Kreft et al. (2018) (violet bar).

### B. *Experiment 2:* Auditory enhancement under forward masking

#### 1. Stimuli

Stimuli were similar to Experiment 1, except that the 500-ms precursor and 100-ms-duration masker were followed by a 20-ms gap and then a 20-ms-duration probe tone at the target frequency of 4000 Hz. Precursors, maskers, and the probe were gated on/off with 10-ms cos^2^ ramps. Five stimulus conditions were simulated (Kreft and Oxenham, 2019; Fig 9):

**Figure 9:**
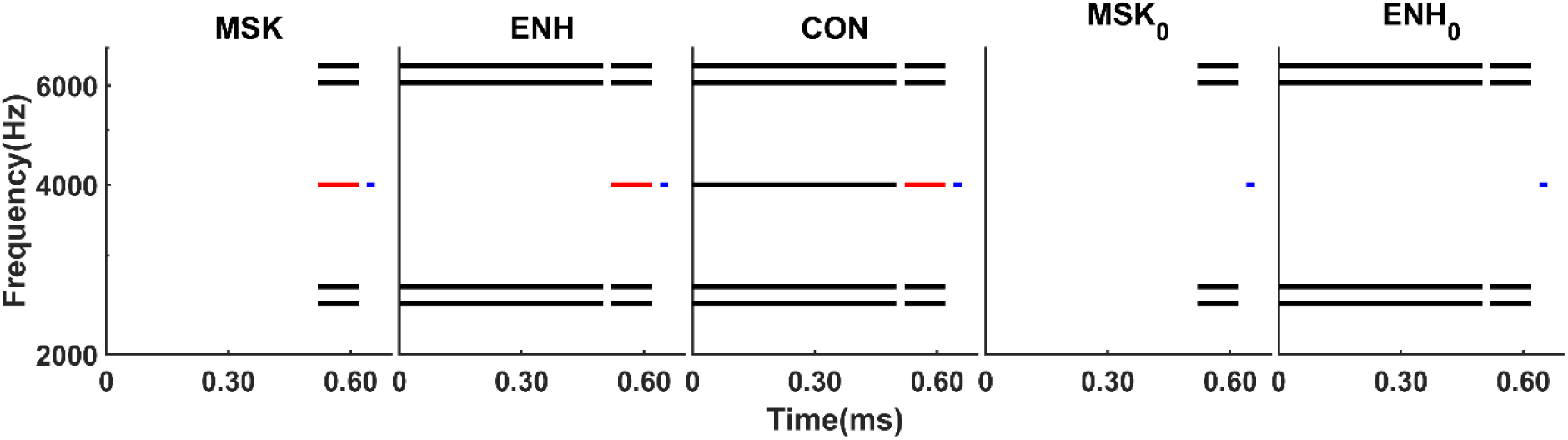
Schematic diagrams of the AE under forward-masking conditions used in Kreft and Oxenham (2019). In the masker-only condition (MSK), the masker is followed by a probe (blue). The enhanced condition (ENH) includes a precursor without the target component, expected to enhance the target component of the masker (red), thereby increasing the masking effect on the final probe (blue). The control condition (CON) also includes a precursor, but with the target component present, eliminating expected enhancement despite the precursor’s presence. The MSK_0_ and ENH_0_ conditions are identical to the MSK and ENH conditions, respectively, except that the masker lacks the target component. The simulations employed a two-interval, two-alternative forced-choice task, with only the intervals containing the probe (in red) shown here.

MSK = Masker that included target component (Baseline)

ENH= Added precursor without target component (Enhanced condition)

CON = Control; Precursor contained target component (No Enhancement)

MSK_0_ = Masker without target component

ENH_0_ = Precursor and masker both lacked target component

The latter two conditions (MSK_0_ and ENH_0_) were to estimate the effects of the masker or precursor duration.

For simulations of listeners with HI, stimuli were always 85 dB SPL/component, as in Kreft and Oxenham (2019). For simulations of listeners with NH, simulations were done in three level configurations:

NH85: SPL matched to HI (85 dB SPL/component),

NHSL: Sensation level (SL) matched to average HI SL (27.4 SPL/component), and

NMNH: SPL (85 dB SPL/component) and SL matched to the average HI listener using threshold equalizing noise (TEN, 50 dB SPL). TEN noise was gated on 100-ms prior to the start of the first interval and gated off 100-ms after the end of the last interval in each trial.

#### 2. Anticipated Result

We hypothesized that AE in the forward-masking paradigm is explained by differences in the effect of the MOC on cochlear gain, as follows: In the ENH condition, even though the precursor lacks a target-frequency component, the target-frequency channel would be excited by the beating between the low-frequency masker components. The IC response to the beating would drive the MOC and reduce gain in the target frequency channel, elevating the probe-detection threshold. In the MSK condition, the lack of a precursor would result in little gain reduction in the target-frequency channel. Interestingly, for the CON condition, the presence of the target-frequency component in the precursor would result in capture of AN responses in the target-frequency channel (Carney, 2024). Capture would overwhelm the response to the beating of the low-frequency precursor components; the captured response would be less effective in exciting the IC model, resulting in less gain reduction via the MOC. Thus, the elevation of the probe-detection threshold would be reduced in the CON condition as compared to the ENH condition. This mechanism for AE would be level-dependent, because high-SPL stimuli are required for the beating of low-frequency precursor components to effectively stimulate the target-frequency channel. Reduced cochlear gain throughout the precursor, masker, and probe, due to hearing loss or due to both compression and the MOC reflex in response to the TEN noise (NMNH), would minimize the effect of these gain changes, consistent with the lack of AE in these conditions.

#### 3. Procedure

Model thresholds for probe detection were estimated using a two-interval, forced-choice, method-of-constant-stimuli paradigm with an inter-stimulus interval (ISI) of 500-ms. A total of 50 trials were simulated for each condition. Odd trials had the probe tone in the first interval, whereas even trials had the probe tone in the second interval.

Probe level was varied from 10 to 100 dB SPL, with 5-dB increments to estimate detection thresholds for NH and HI models. For NMNH, the TEN noise level was determined by setting the probe level to the average threshold for listeners with HI and adapting the TEN level to just mask the probe for the paired NH listener. The TEN level was thus adjusted to 50 dB SPL, which was similar the mean TEN noise level used in Kreft and Oxenham (2019). Assuming that listeners depended on responses in frequency channels tuned near the probe frequency, the decision variable was based on rates summed across seven IC-BE model neurons with CFs of 3.6, 3.7, 3.8, 4, 4.2, 4.3, and 4.5 kHz (Brennan et al. 2023). The interval selected as having the probe was the one with the maximum instantaneous model-IC rate in a time window that included the final 25 ms of the masker and the entire probe response (Fig. 10). This time window was chosen because listeners were assumed to detect some change at the end of the stimulus, rather than simply responding to a longer-duration sound (Brennan et al., 2023).

**Figure 10:**
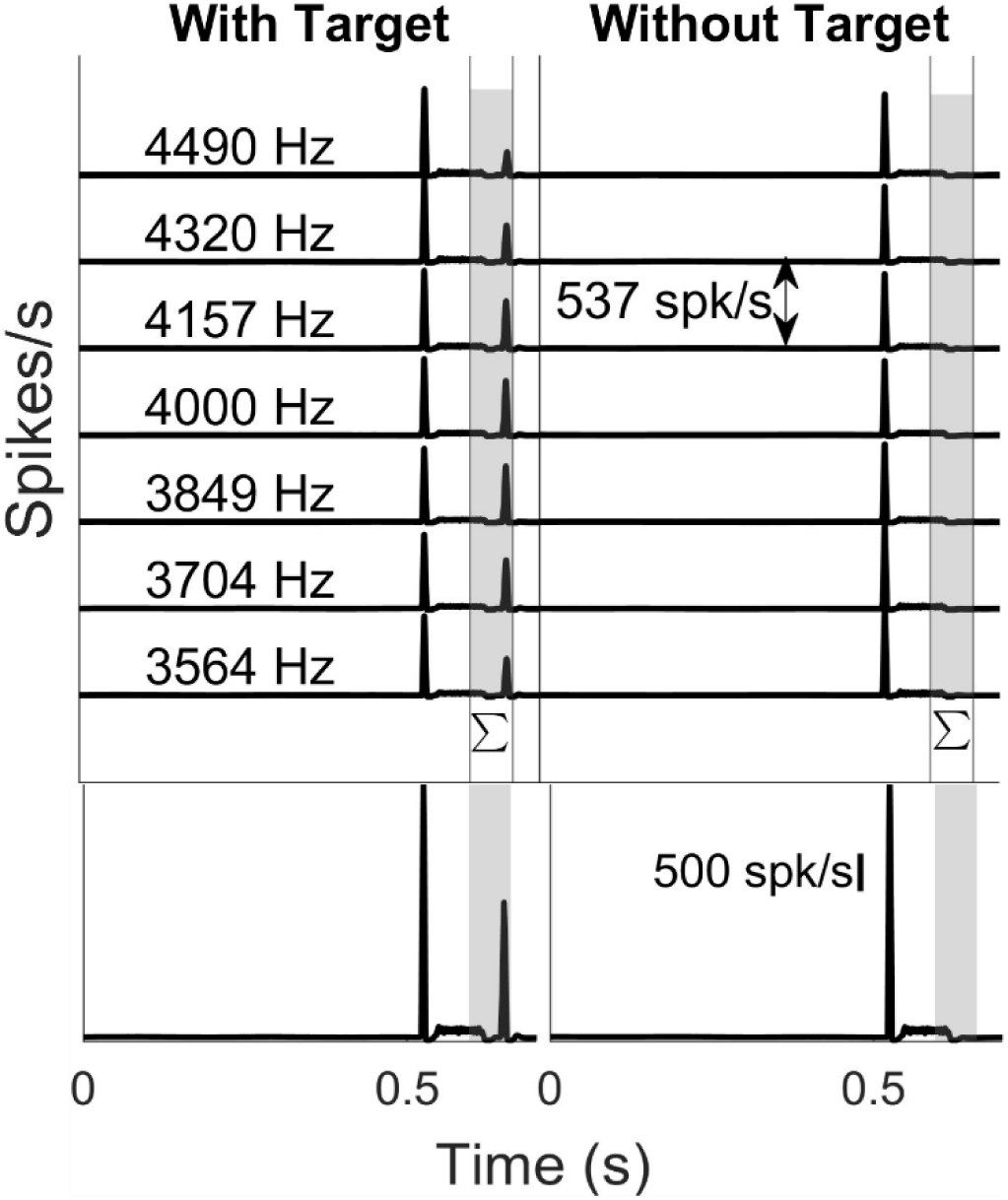
The profile across CF of instantaneous rates of model-IC responses for AE under the forward masking paradigm, with and without the target. The maximum rate was computed over the time window that included the final 25 ms of the masker and extended through the probe response across the CFs (gray shaded area). The target level was set 5 dB above detection threshold for this illustration.

The percent correct was computed for each probe level, and a logistic function was then fit to the percent-correct vs. level curve. Threshold was estimated as the level for which the curve intersected the 70.7%-correct point. This process was repeated for each stimulus type and participant group.

#### 4. Results

The effect of efferents on the AN-model responses for NH listeners with SPL matched to HI (85 dB SPL/component) in the ENH condition is shown in Fig. 11. For the AN models with and without efferents, in the AN channel tuned near the target frequency neural fluctuations driven by the beating (∼177 Hz) between the low-frequency masker components were observed in the ENH condition during the precursor, because the target frequency was absent and the stimulus level was high enough to elicit off-CF excitation. For CON, in which the target frequency was present during the precursor, capture of the AN response by the target component dominated and the response to the target overwhelmed the fluctuation in the responses. However, no difference was observed during the probe window for the response to the MSK, ENH, and CON conditions in the responses of the AN model without efferents (Fig. 11, top half). But in the model with efferents (Fig. 11, bottom half), differences in model responses in the probe window (gray shaded area) were observed across conditions. The cochlear gain in the model was affected by compression, suppression, and by the cochlear-gain factor from the output of the model MOC stage. The combined effects of these factors can be observed in the time constants of the cochlear filters (Fig. 12). In the model with efferents, and in the NH85 configuration (for which AE is observed), the time constant (and thus the cochlear gain) was highest during the probe for the MSK condition (blue) followed by the ENH (red) and CON (yellow) conditions (Fig. 12, top row). Note that in the model without efferents, there were no differences in the peripheral-filter time constant across conditions (not shown).

**Figure 11:**
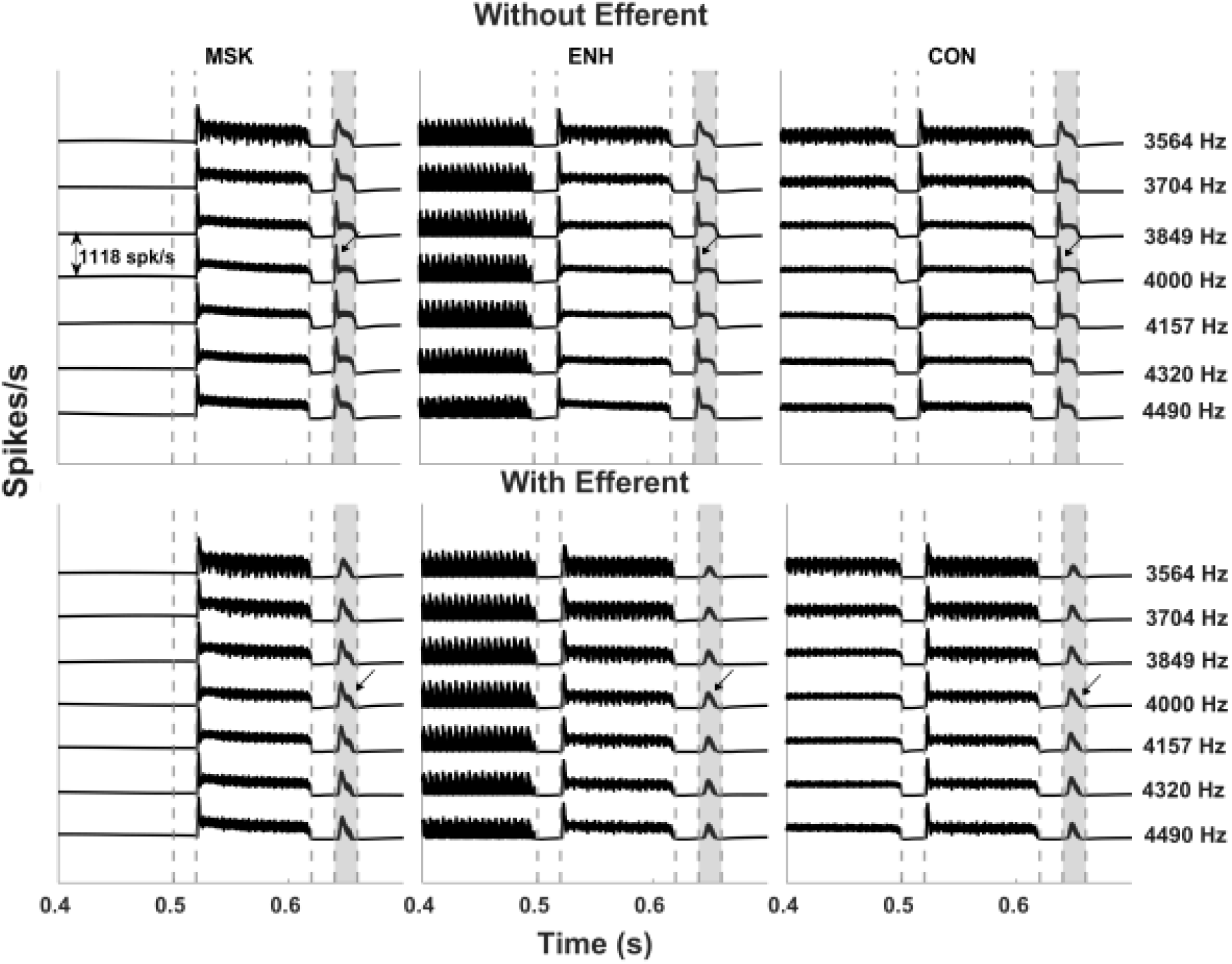
Efferent gain control results in differences in model responses at the level of AN for AE under forward masking. Response profiles for the frequency channel around the target frequency channel (indicated by arrows) are shown as function of time. The dashed vertical lines in gray indicate the precursor offset, masker onset, masker offset, probe onset, and probe offset, from left to right. The target is set 5 dB above the threshold observed for ENH condition. Responses of the model without efferents (top half) shows little differences in the probe response (gray area) across stimulus conditions for the NH model in quiet (NH85), which has SPL matched to that for listeners with HI. Responses of the model with efferents show lower responses for probe (gray area) for ENH compared to MSK and CON, explaining increased model detection threshold for ENH condition.

**Figure 12:**
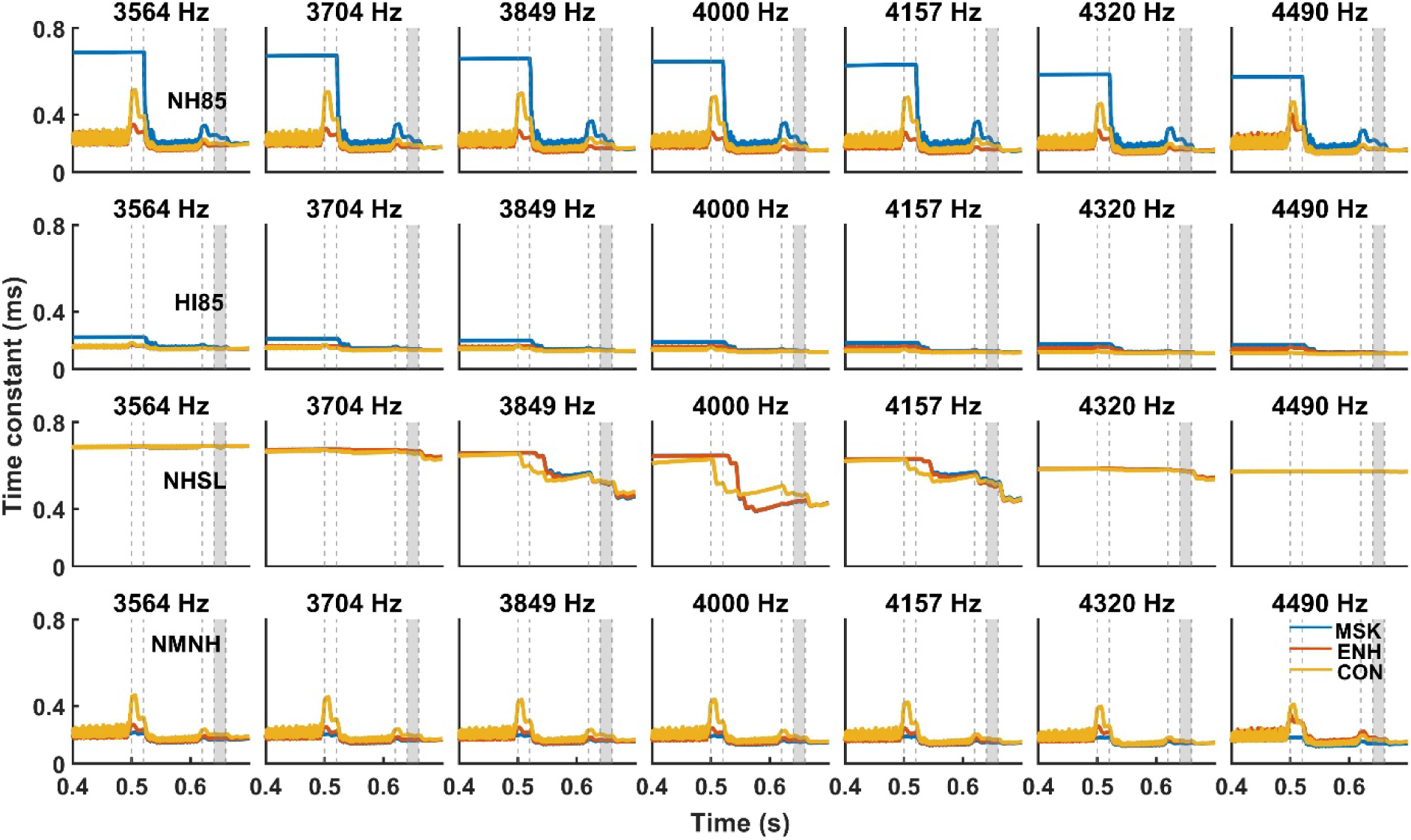
Changes in the time constant of the peripheral filter (the C1 filter in Zilany et al., 2014) in the narrow frequency range around the target frequency explains the enhancement effect reported by psychoacoustic studies for AE under forward masking. The dynamics of the time constant, which is proportion to cochlear gain, are shown for MSK (blue), ENH (red) and CON (yellow) conditions. The target was set 5 dB above the threshold observed for the ENH condition. The dashed vertical lines in gray indicate the precursor offset, masker onset, masker offset, probe onset, and probe offset, from left to right. Differences in the time constant, and thus cochlear gain, between MSK and ENH in the NH85 configuration during the probe window (gray shaded area) explain AE in the model results.

The trends in the AN model responses were reflected in the IC responses (Fig. 13), in which the differences among conditions were further pronounced. For the model with efferent gain control, the probe response was higher in the MSK condition (Fig. 13, bottom row in each panel) than in the ENH condition (Fig. 13, top row in each panel), making probe detection more challenging in the ENH condition. Consequently, thresholds were higher for ENH compared to MSK for NH listeners when SPL was matched to HI (NH85, Fig. 13A). Although there is more cochlear gain for NHSL than NH85 (Fig. 12), enhancement was not observed in NHSL because the stimulus level (27.4 dB SPL/component) was not sufficient to evoke substantial response differences between the ENH and MSK conditions (i.e., an effect of the target component on responses to off-CF excitation due to the beating of low-frequency components) (Fig. 13C), unlike in the NH85 condition (85 dB SPL/component) (Fig. 13A). In all other conditions, for the model with or without efferents, the difference in probe response between MSK and ENH was less pronounced (Fig. 13B HI; Fig. 13D NMHL).

**Figure 13:**
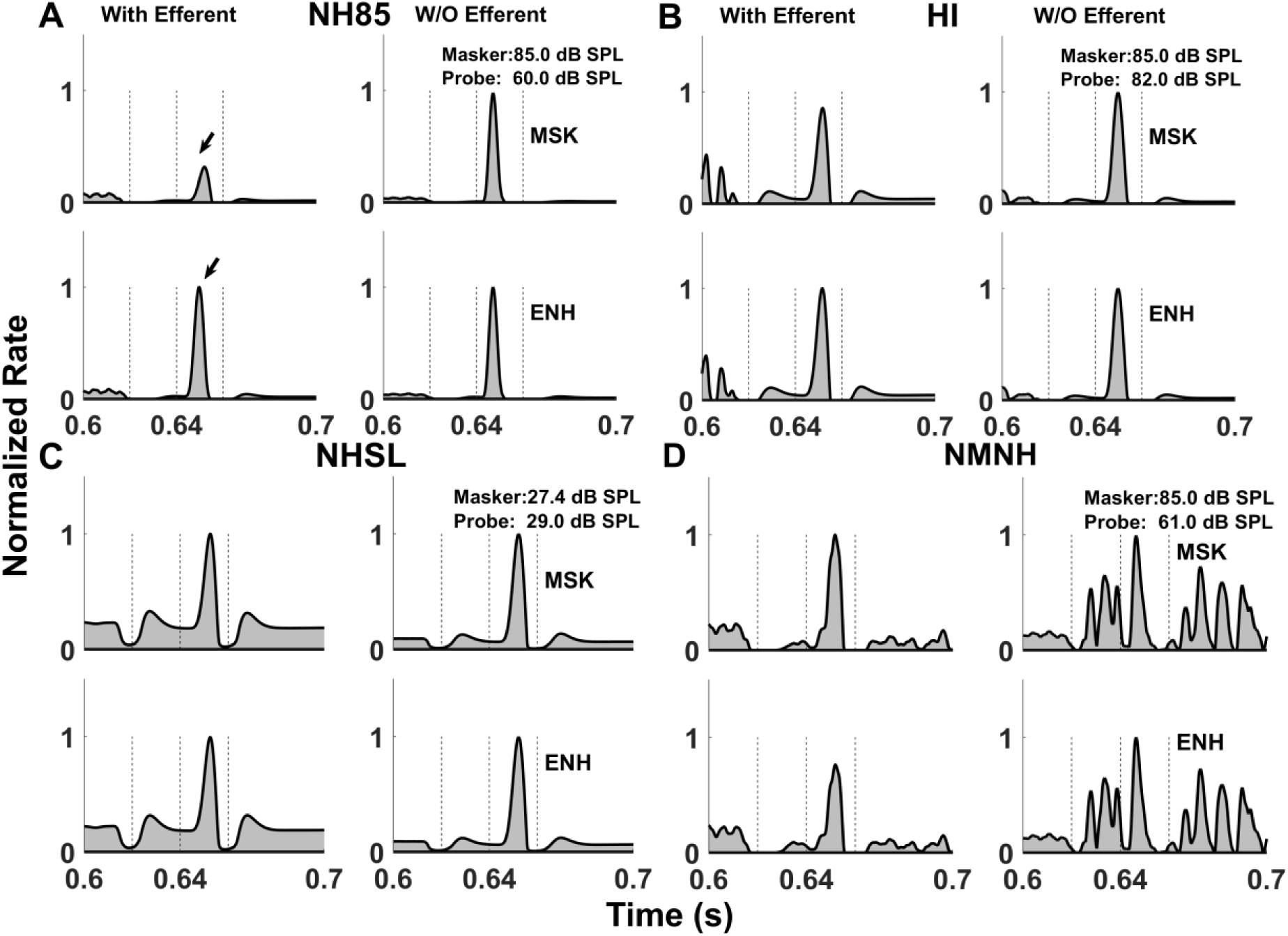
AE under forward masking occurs only in NH85 condition, and only in models with efferent gain control (arrow). Responses of models with and without efferents to the probe, summed across CFs near the probe frequency, as a function of time. For each configuration, responses were normalized by maximum response. The target response is shown for MSK (bottom) and ENH (top) for each condition. Probe levels were set 5 dB above threshold for the ENH condition. A) NH85: NH group with stimuli presented at a similar SPL as the HI group. B) HI group response. C) NHSL: NH group with stimuli presented at the same SL as for the HI listeners. D) NMNH: NH group with stimuli presented in a background of TEN to simulate the hearing loss of the matched HI listener, with masker and enhancer presented at the same SPL as for the HI listeners. The dashed three vertical lines indicate the masker offset, probe onset and probe offset respectively.

The thresholds for the model with efferent feedback (orange circles) closely matched the experimental data (violet squares), whereas the model without efferents (unfilled gray circles) remained similar across most conditions (Fig. 14). Because cochlear gain was constant in the model without efferents, variations in conditions had no impact on threshold, unlike in the model with efferents.

**Figure 14:**
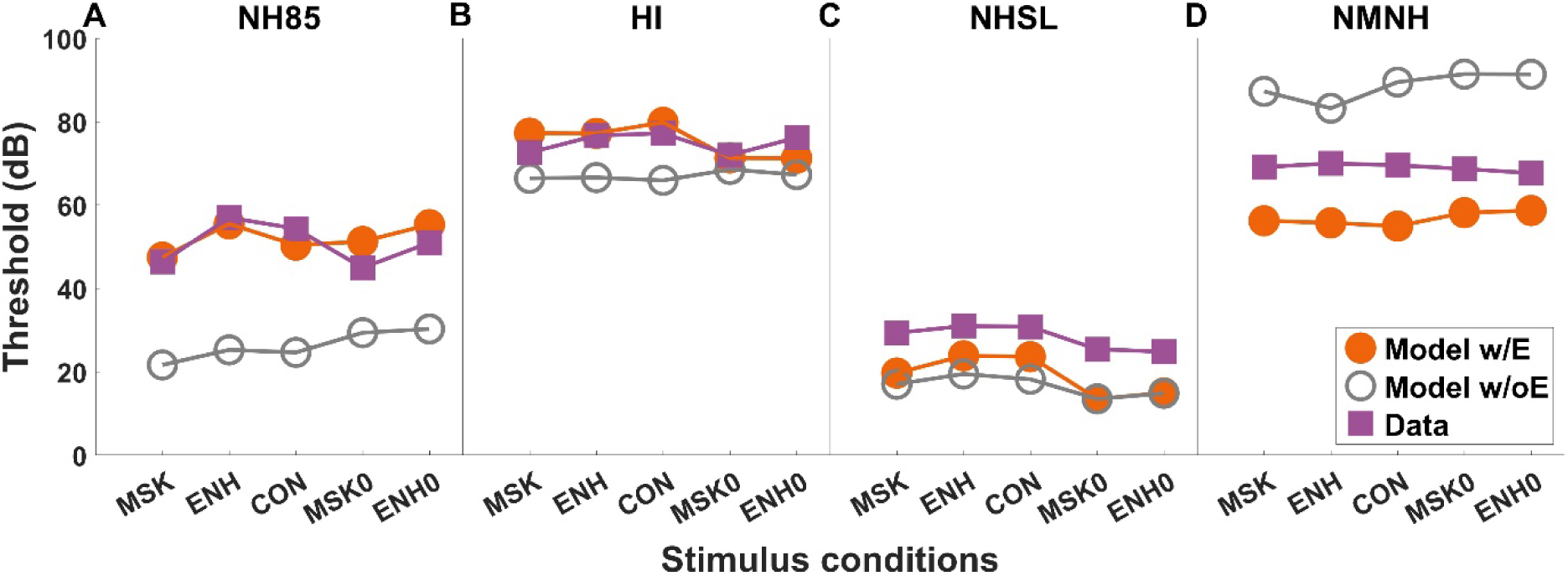
Thresholds as a function of stimulus conditions for a target frequency of 4000 Hz across different configurations used to explore AE under forward masking: (A) NH85 (B) HI (C) NHSL (D) NMNH. Data from Kreft and Oxenham (2019) are shown as violet squares, model with efferent (w/E) as orange filled circles, and model without efferent (wo/E) as gray unfilled circles.

Following the approach of Kreft and Oxenham et al. (2019), we assessed the effect of duration by measuring forward-masked thresholds for the probe under two conditions: with the masker alone, and with the masker plus precursor—neither containing the target component. The difference in thresholds between these two conditions estimates the duration effect in the absence of the target (ENH_0_-MSK_0_). The change in probe threshold attributable to increased masker duration was calculated as (ENH-MSK) - (ENH_0_ - MSK_0_) (Fig. 15). In the HI model with efferent feedback, this effect was minimal; in contrast, in the NH model with efferents demonstrated, the effect of duration closely matched the experimental data for both the SPL-matched (NH85) and SL-matched (NHSL) configuration, while remaining low in the NMNH configuration.

**Figure 15:**
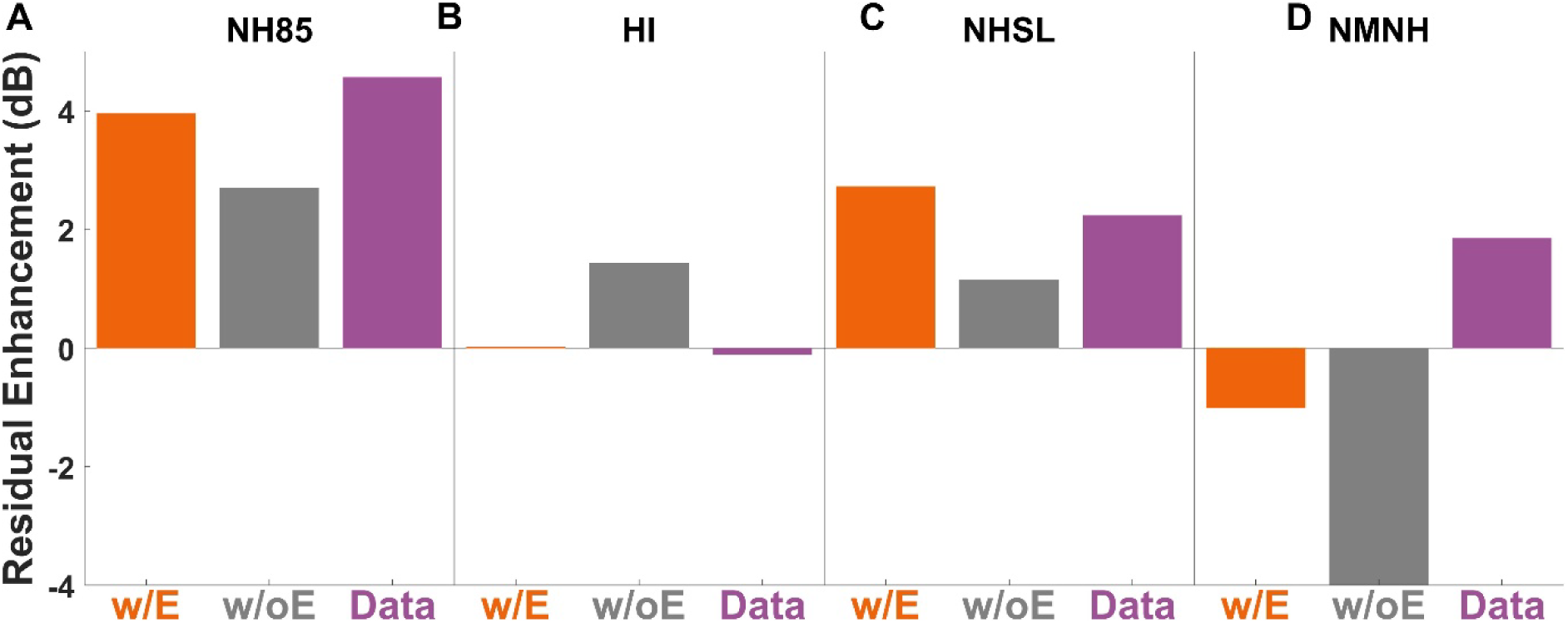
Residual enhancement effect (in dB) measured by (ENH-MSK)-(ENH_0_-MSK_0_) for a target frequency of 4000 Hz across different configurations of AE under forward masking: A) NH85 (B) HI (C) NHSL (D) NMNH. Results are shown for the model with efferent (orange bars), model without efferent (gray bars), and data from Kreft and Oxenham (2019) (violet bars).

## IV. Discussion

As evidenced by previous psychophysical studies in listeners with NH or HI (Kreft et al. 2018; Kreft and Oxenham, 2019), AE may occur through distinct yet potentially overlapping mechanisms under simultaneous- and forward-masking paradigms. Our model incorporating MOC efferent gain control exhibited AE effects that were consistent with psychoacoustic studies for both NH and HI listeners. In contrast, a model lacking efferent gain control failed to capture AE effects. In the first experiment, AE under simultaneous masking, enhancement was observed across all configurations, consistent with experimental data (Kreft et al. 2018).

In the NH model, enhancement varied with stimulus level, similar to the trends in psychoacoustic studies (Kreft and Oxenham, 2019). Additionally, a residual enhancement effect was present in the NH model when the sound level was matched to that of the HI condition (NH85), but this effect did not appear in other NH configurations. The residual enhancement at high SPL was attributed to the response of the target-frequency channel to the beating of low-frequency precursor components. The IC response to the beating would reduce the cochlear gain unlike in other conditions for which the cochlear gain was already reduced, either due to hearing loss, or to both compression and the MOC reflex in response to the TEN noise (NMNH), or to low sound level (NHSL).

The enhancement in simultaneous masking for the HI model was due to gain changes in off-target channels. However, in the second experiment, AE under forward masking, no enhancement was observed in the HI model. In forward masking, the models focused on the gain in the target channel, which is critical for detection of a low-level probe tone. In the HI model, the target-channel gain was similar between enhanced and unenhanced conditions, so no AE effect was observed.

These findings supported the hypothesis that MOC efferent gain control plays a role in AE. The decision variables used to model the two tasks differed. In AE under simultaneous masking (Fig. 3), the decision variable was derived from a frequency profile, emphasizing the target frequency more than the off-target frequencies. The stimulus consisted of a precursor followed by a masker containing both the target and off-target components, potentially requiring the listener to selectively attend to the target amidst competing spectral elements. In contrast, under forward masking, the decision variable was narrowly tuned to frequencies around the target (Fig. 8). Since the probe tone was presented alone following the masker, listeners likely rely on a narrow frequency range, unlike in simultaneous masking where off-target frequency channels were also involved. For listeners with hearing loss, enhancement was observed under simultaneous masking (Fig. 6B) and not under forward masking (Fig. 11B), likely because the simultaneous condition utilized a broader range of frequency channels compared to the narrowly focused processing in forward masking. The involvement of a wider range of channels might provide more opportunity to construct a spectral profile, even if some channels were functioning suboptimally.

Further insights into AE mechanisms come from studies on individuals with hearing loss and cochlear implants (CIs). If AE relies on the normal functioning of the cochlea, individuals with sensorineural hearing loss (SNHL) should exhibit diminished enhancement due to outer-hair-cell dysfunction (Ruggero and Rich, 1991; Oxenham and Plack, 1997). In line with this, AE appears to be reduced in listeners with SNHL, particularly under forward-masking conditions (Thibodeau, 1991; Kreft and Oxenham, 2017).

CI users exhibit AE under simultaneous masking but not under forward masking (Kreft and Oxenham, 2017). This result is typically interpreted as suggesting that AE may arise from two distinct mechanisms: one that is preserved in CI users, potentially central in nature, and another that is absent in CI users, likely reliant on peripheral auditory processing (Kreft and Oxenham, 2017; Wang et al., 2015, 2016). However, the MOC system projects directly to AN fibers below the inner hair cells, in addition to the better-known projection of the lateral efferent system to the afferent fibers (Hua et al. 2021). Thus, descending central projections could influence neural representations at the level of the AN, potentially influencing phenomena such as AE and forward masking (Maxwell et al., 2024), even when the auditory periphery is bypassed by CIs. Overall, AE may involve both central and peripheral mechanisms, with contributions from cochlear gain control and efferent modulation, which refers to the changes in neural responses due to the MOC influence on the outer hair cells and cochlear gain.

Beim et al. (2015) proposes that MOC efferent activation by a precursor without a target frequency could lead to a frequency-specific reduction in basilar membrane gain for the masker components. Additionally, given the longitudinal coupling within the cochlea, the gain reduction might, in turn, reduce suppression by the masker of the basilar membrane response to the target. Such a decrease in suppression was expected to manifest as an increase in the stimulus-frequency otoacoustic emission (SFOAE) magnitude at the target frequency when the precursor is present, compared to the condition without the precursor. Therefore they hypothesized that if efferent gain control played a role in AE under simultaneous masking, the gain, and thus SFOAE magnitude, at the target frequency should either increase, or remain unchanged while the gain at the flanking masker components was reduced (Beim et al., 2015). Our model showed reduced gain at masker frequencies but no change in gain at the target frequency (Fig. 5), supporting their hypothesized frequency-specific MOC effects (Beim et al., 2015), but not the hypothesized release-from-suppression mechanism. Note that their hypothetical framework was based on the energy-driven inputs to the MOC system, and did not include the descending, fluctuation-sensitive pathway from the inferior colliculus.

Beim et al.’s (2015) SFOEA measurements did not support either of their hypotheses; they do not report significant changes in SFOAE magnitudes across MSK and ENH conditions at either target or masker frequencies. Even when avoiding middle-ear muscle reflexes by using lower-level, narrowband inharmonic complexes (which still produce psychophysical enhancement), SFOAE changes remain undetectable. However, the absence of measurable changes in the SFOEA does not rule out efferent involvement because, as Beim et al. (2015) pointed out, the expected cochlear gain changes could have been smaller than 1 dB, below the detection threshold for SFOAEs, and the use of 2-sec silent intervals may not have allowed full recovery from MOC activation, especially at higher frequencies, where MOC effects decay more slowly, potentially minimizing differences across conditions. Goodman et al. (2025) suggest that the minimum detectable change could be a statistically rigorous tool for interpreting individual MOC reflex results, particularly when responses are non-significant, because it may help to distinguish whether a lack of significance is due to a genuinely weak or absent MOC reflex, or simply a poor signal-to-noise ratio (SNR). Given the measurement challenges, computational modeling provides a valuable complementary approach for demonstrating and understanding efferent influences. In our model simulations of AE under simultaneous masking, the cochlear-gain changes (Fig. 5) averaged over the entire masker stimulus duration were 2.6, 3.2, 0.3, 2.8, and 2.1 dB for channels tuned to the 2.4, 2.6, 4, 6.0, and 6.4 kHz components, respectively. Except at the target frequency, these gain changes should be large enough to be detected with SFOAEs, although our estimates were not affected by phase shifts, which could affect the SFOAE estimates. Large gain changes can be elicited by tones; for example, gain is reduced by approximately 20 dB with ipsilateral MOC elicitors in a psychophysical study (Drga et al., 2016) and in physiological studies of AN responses that used electrical shocks or acoustic elicitors to activate the efferents (e.g. Guinan and Gifford, 1988; Warren and Liberman, 1989).

Experimental factors such as probe level (threshold vs. suprathreshold), use of suppressor tones, and stimulus configuration can significantly influence SFOAEs and how they reflect cochlear sensitivity. For example, using SFOAEs, Keefe et al. (2009) did not observe an overshoot-like difference between the short- and long-delay conditions for a tone-in-noise detection task, whereas Walsh et al. (2010) did, highlighting the role of methodology in SFOAE results.

A previous electroencephalogram (EEG) study by Mehta et al. (2021) shows strong correlates of perceptual enhancement in the low-frequency auditory steady-state response (ASSR) (∼40 Hz), which is believed to reflect cortical activity (Plourde et al., 2006; Herdman et al., 2002). However, no corresponding enhancement is observed in the higher-frequency ASSR (100–200 Hz), typically linked to subcortical sources (Bidelman et al., 2018). This absence does not necessarily rule out AE at subcortical levels, as (i) some participants exhibit increased 100- and 200-Hz ASSR amplitudes (Mehta et al., 2021), (ii) subcortical activity may not be reliably detected by surface EEG electrodes, possibly because subcortical neurons involved in enhancement do not effectively entrain to the stimulus envelope (Mehta et al., 2021), and (iii) a neuronal correlate for AE has been observed in the IC of awake marmoset (Nelson and Young, 2010).

Our model simulations support the hypothesis that efferent gain control contributes to AE, but additional mechanisms may also contribute to the observed effects. It should be noted that our model is poorly constrained due to limited physiological data on the magnitude and temporal dynamics of efferent gain control. As an example, the differences in probe-detection thresholds for the models with and without efferents (Fig. 14, orange vs. grey symbols) were relatively large (∼25-35 dB), but there is little information in the physiological literature to inform the constraint of this effect size. In the model, the cochlear-gain factor varied between 0 and 1, and the maximum change in cochlear gain was only constrained by the amount of available gain in each frequency channel. The available gain depended on both the CF and the degree of hearing loss for that frequency, which was adjusted based on the average audiogram of the subjects within each group.

Enhancement has also been observed with long precursor-masker gaps of up to 1 s (Viemeister, 1980; Feng and Oxenham, 2015). A recent physiological study of forward masking showed that a 1-s delay between masker and probe is insufficient for recovery from masking (Agarwalla et al., 2025). The model with efferents captures the physiological trend in recovery for delays up to 500 ms, but fails for longer delays. However, the model did not include slow efferent effects which can last several seconds (Cooper and Guinan, 2003; Backus and Guinan, 2006). These slow effects may be critical for explaining enhancement observed at longer delays.

Future studies using animal models and employing behavioral paradigms could provide further insight into the neural locus of AE, helping to determine whether it is mediated by feedforward mechanisms, efferent cochlear gain control, or an interaction between the two. Experimental approaches such as cortical deactivation through cooling, optogenetic silencing, or pharmacological manipulations may help clarify these mechanisms. Further investigations, particularly those integrating physiological recordings with behavioral paradigms, will be essential in determining the source of AE.

Summerfield and Assmann (1989) shows that AE improves listeners’ ability to segregate and identify concurrent vowels. The benefit is attributed to enhanced perceptual grouping and auditory object segregation—processes essential for speech comprehension in acoustically complex environments. If AE facilitates the perception of concurrent vowels, and if efferent activity underlies or supports AE, then efferent mechanisms are likely to contribute to speech perception, especially under challenging listening conditions.

## ACKNOWLEDGEMENTS

We thank Heather Kreft for sharing the TEN noise thresholds and Daniel R. Guest for helpful comments. The work was supported by NIH Grants R01-DC010813 (LHC, SA), R01-DC009838 (AF), and F32-DC022782 (AF).

## Author Declarations

The authors have no conflicts of interest to disclose.

## Data Availability

The code used for the simulations described here will be made available at https://osf.io/ufqpc/overview

## Notes

### Competing Interest Statement

The authors have declared no competing interest.

